# Requirement of cholesterol for calcium-dependent vesicle fusion by stabilizing synaptotagmin-1-induced membrane bending

**DOI:** 10.1101/2022.02.03.478933

**Authors:** Houda Yasmine Ali Moussa, Kyung Chul Shin, Janarthanan Ponraj, Soo Jin Kim, Je-Kyung Ryu, Said Mansour, Yongsoo Park

**Author notes:** Corresponding authors; Dr. Yongsoo Park, Neurological Disorders Research Center, Qatar Biomedical Research Institute (QBRI), Hamad Bin Khalifa University (HBKU), Qatar Foundation, Doha, Qatar. These authors contributed equally to this work (Houda Yasmine Ali Moussa, Yongsoo Park). **Competing Interests:** The authors declare no competing interests.

## Abstract

Cholesterol is essential for neuronal activity and function. Cholesterol depletion in the plasma membrane impairs synaptic transmission. However, the molecular mechanisms by which cholesterol deficiency leads to defects in vesicle fusion remain poorly understood. Here we show that cholesterol is required for Ca^2+^-dependent native vesicle fusion using the *in-vitro* reconstitution of fusion and amperometry to monitor exocytosis in chromaffin cells. Purified native vesicles were crucial for the reconstitution of physiological Ca^2+^-dependent fusion, whereas vesicle-mimicking liposomes failed to reproduce the cholesterol effect. Intriguingly, cholesterol had no effect on membrane binding of synaptotagmin-1, a Ca^2+^ sensor for ultrafast fusion. Cholesterol stabilizes local membrane bending induced by synaptotagmin-1, thereby lowering the energy barrier for Ca^2+^-dependent fusion to occur. Our data provide evidence that cholesterol depletion abolishes Ca^2+^-dependent vesicle fusion by disrupting synaptotagmin-1-induced membrane bending, and suggests that cholesterol is an important lipid regulator for Ca^2+^-dependent fusion.

---

Cholesterol is a major component in cell membrane bilayers, and is essential for membrane structure and fluidity. Brain is the most cholesterol-enriched organ and a human brain contains about 20–25% of the body’s cholesterol^1, 2^; this high density suggests that cholesterol has a critical function in the brain. AgeLJrelated cholesterol reduction in the frontal and temporal cortices^3^ results in loss of synaptic contacts, changes in neuronal morphology, and reduced synaptic plasticity^4^. Cholesterol is associated with neurogenesis, neurodevelopment, and synaptogenesis^5, 6^. Age□related cholesterol deficiency in the plasma membrane leads to deficits in synaptic plasticity in mouse hippocampal neurons^7^, and cholesterol depletion by methyl-β-cyclodextrin (MCD) from the plasma membrane impairs neurotransmission and neuronal activity, and thereby leads to synapse degeneration^8, 9^.

The plasma membrane contains ∼80% of total cellular cholesterol^10^. Cholesterol is capable of clustering syntaxin-1A^11^ so that soluble *N*-ethylmaleimide-sensitive factor attachment protein receptor (SNARE) proteins become concentrated in cholesterol-enriched domains in the plasma membrane^12^. Depletion and reduction of cholesterol level in the plasma membrane inhibit Ca^2+^-dependent exocytosis of large dense-core vesicles (LDCVs)^13^ and cortical secretory vesicles from sea urchins^14^, as well as synaptic vesicles in hippocampal neurons^8, 15^, cortical synaptosomes^16^, ribbon synapses^17^, and motor nerve terminals^18^. However, the molecular mechanisms by which cholesterol deficiency disrupts synaptic transmission and induces neurodegeneration remain elusive and controversial.

Exocytosis is the process of neurotransmitter release through merging two lipid bilayers. It is mediated by SNARE proteins^19, 20^. Neuronal SNARE proteins consist of Q-SNARE in the plasma membrane (syntaxin-1 and SNAP-25) and R-SNARE in the vesicle membrane (synaptobrevin-2 or vesicle-associated membrane protein-2 (VAMP-2))^19^. Synaptotagmin-1 is responsible for ultrafast Ca^2+^- dependent exocytosis. Synaptotagmin-1 contains two tandem C2-domains that coordinates Ca^2+^ binding; the Ca^2+^-bound C2 domain can be inserted to negatively charged anionic phospholipids by electrostatic interaction^20^. In spite of intense investigation of synaptotagmin-1, the molecular mechanisms by which synaptotagmin-1 mediates Ca^2+^-dependent vesicle are still under debate^21^.

Here we propose novel mechanisms by which cholesterol in the plasma membrane mediates synaptotagmin-1-induced vesicle fusion. We used amperometry to monitor exocytosis in real time, and performed *in-vitro* reconstitution of vesicle fusion including large dense-core vesicles (LDCVs) and synaptic vesicles (SVs), and showed that cholesterol is required for Ca^2+^-dependent vesicle fusion. Importantly, vesicle-mimicking liposomes fail to reproduce the cholesterol effect, but purified native vesicles, i.e., LDCVs and SVs, are crucial for the reconstitution of physiological Ca^2+^-dependent fusion. Membrane binding of synaptotagmin-1 occurs regardless of cholesterol. Transmission electron microscope (TEM) revealed that cholesterol stabilizes membrane bending and curvature induced by the insertion of synaptotagmin-1, and therefore the membrane bending energy can lower the energy barrier to fusion.

## Results

### Cholesterol depletion causes deficits in exocytosis

We investigated whether local membrane curvature is observable *in-vivo* using primary chromaffin cells. Using electron microscopy (**Fig. 1a-f**) we observed the invagination of the plasma membrane into LDCVs, suggesting that the plasma membrane could be already curved before Ca^2+^ triggering. Cholesterol induces spontaneous membrane curvature and bending^22, 23^, thereby promoting highly curved membrane intermediate structures in membrane fusion^24, 25^. Moreover, high-curvature membrane domains such as caveolae are heavily enriched in cholesterol, which is essential for membrane invaginations^22, 26^. Cholesterol depletion by methyl-β-cyclodextrin (MCD) or nystatin disrupts membrane invaginations and leads to a flattening of the curved caveolae membrane structure^27, 28^.

**Figure 1.**
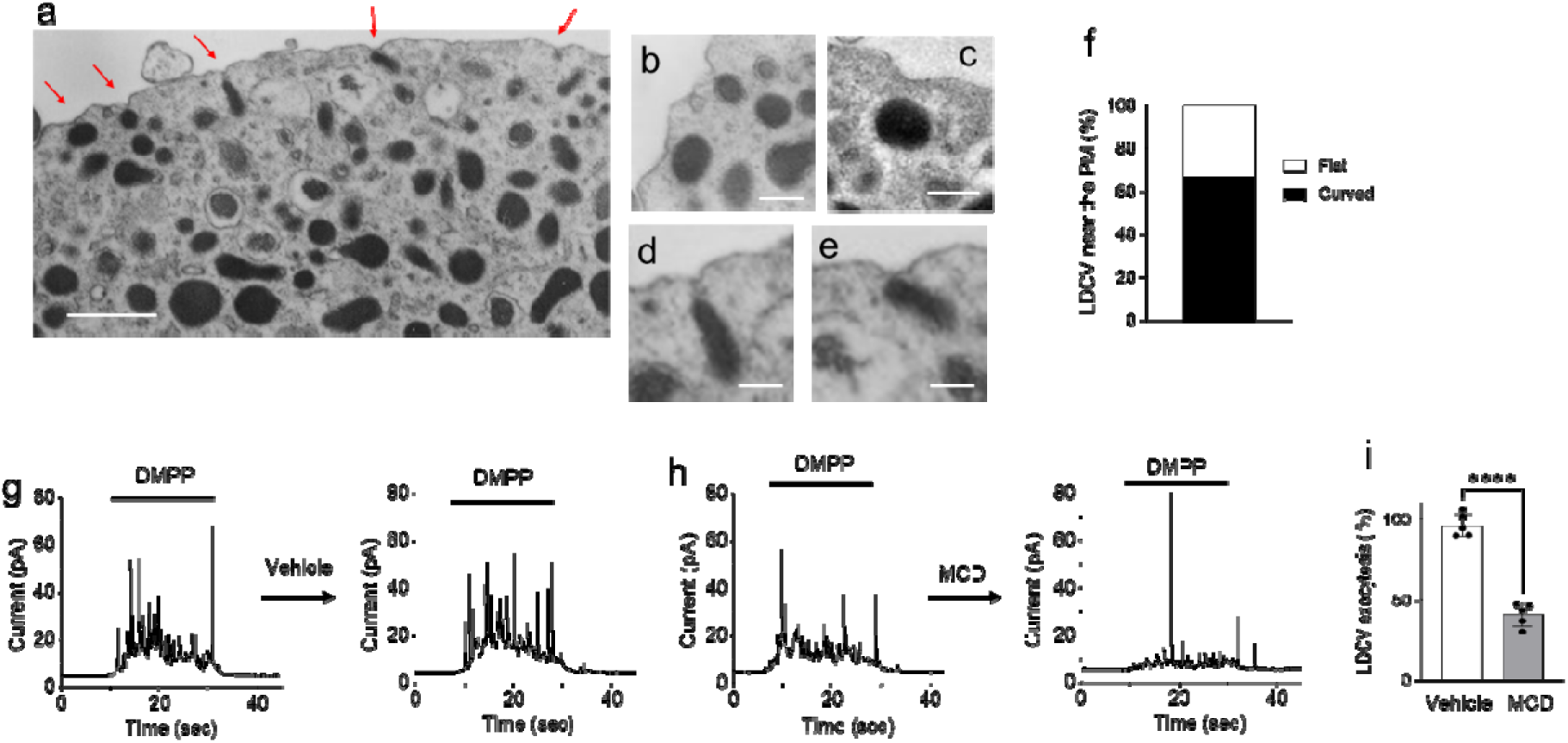
Cholesterol depletion inhibits LDCV exocytosis in chromaffin cells. (**a**) Transmission electron microscope (TEM) image of chromaffin cells showing invagination of the plasma membrane (PM) into LDCVs. Scale, 500 nm. (**b-e**) Magnified TEM images of LDCVs. Scale, 200 nm. (**f**) LDCVs close to the PM less than 100 nm distance between LDCV and PM were counted; LDCVs near either flat or curved PM is presented as a percentage (total 36 LDCVs proximal to the PM from three independent experiments). (**g-i**) LDCV exocytosis in chromaffin cells measured by amperometry. MCD (10 mM, 2 h, 37°C) reduced LDCV exocytosis of chromaffin cells (**h**). (**g**,**h**) Shown are typical amperometric traces upon DMPP stimulations finally activating voltage-gated calcium channel for 20 sec. Chromaffin cells were treated with either vehicle (**g**) or MCD (**h**) for 2 h after the first DMPP stimulation. (**i**) Amperometric current generated by repetitive stimulation was individually integrated. Relative LDCV exocytosis is presented as a percentage of the first DMPP-induced total catecholamine release. Data are means ± SD from 5 independent experiments. ****, *p* < 0.0001.

Given that the curved and invaginated membranes are enriched with cholesterol^22, 27, 28^ and ∼67% vesicles are proximal to the invaginated plasma membranes (**Fig. 1e**), the membrane invagination to LDCVs we observed is likely to be cholesterol-enriched regions, where SNARE proteins are present^29, 30^ (**Fig. 1a-e**). To test the cholesterol effect on vesicle fusion, we used amperometry to monitor LDCV exocytosis in real time. Indeed, treatment with MCD, which depletes cholesterol from the plasma membrane, inhibited Ca^2+^-dependent LDCV fusion by ∼60% (**Fig. 1g-i**).

### Without cholesterol, no Ca^2+^-dependent vesicle fusion

To further study the molecular mechanisms by which cholesterol deficiency affects vesicle fusion, we applied a reconstitution system of vesicle fusion by using purified native vesicles such as LDCVs and SVs, as reported previously^31, 32, 33^. The plasma membrane-mimicking liposomes (PM-liposomes) contain the stabilized Q-SNARE complex (syntaxin-1A and SNAP-25A in a 1:1 molar ratio^34^)(**Online Methods**).

We first tested the effect of cholesterol on basal fusion without Ca^2+^. LDCV fusion with the PM-liposomes was readily observed (**Fig. 2a**), but the absence of cholesterol in the PM-liposomes reduced LDCV fusion by ∼50% (**Fig. 2a,d**); the PM-liposomes contain either 25% or 0% cholesterol (Chol). The efficiency of the SNARE assembly and SNARE complex formation was examined using the light chain of tetanus toxin (TeNT), a protease specific for free VAMP-2; i.e. VAMP-2 in the assembled SNARE complexes is resistant to the cleavage by TeNT so that TeNT-resistant VAMP-2 represents the ternary SNARE complex formation^31^. After LDCV fusion with the PM-liposomes as a fusion assay in **Fig. 2a**, TeNT was added to cleave free VAMP-2 and quantify VAMP-2 engaged in the SNARE complex formation. Absence of cholesterol (0% Chol) in liposomes slightly reduced the ternary SNARE complex formation compared to liposomes containing cholesterol (25% Chol) (**Fig. 2b**); this result suggests that cholesterol facilitates basal fusion of LDCVs by increasing the efficiency of SNARE assembly. Cholesterol can enhance SNARE-complex formation, by inducing clustering of SNARE proteins in the plasma membrane^11, 12, 30^.

**Figure 2.**
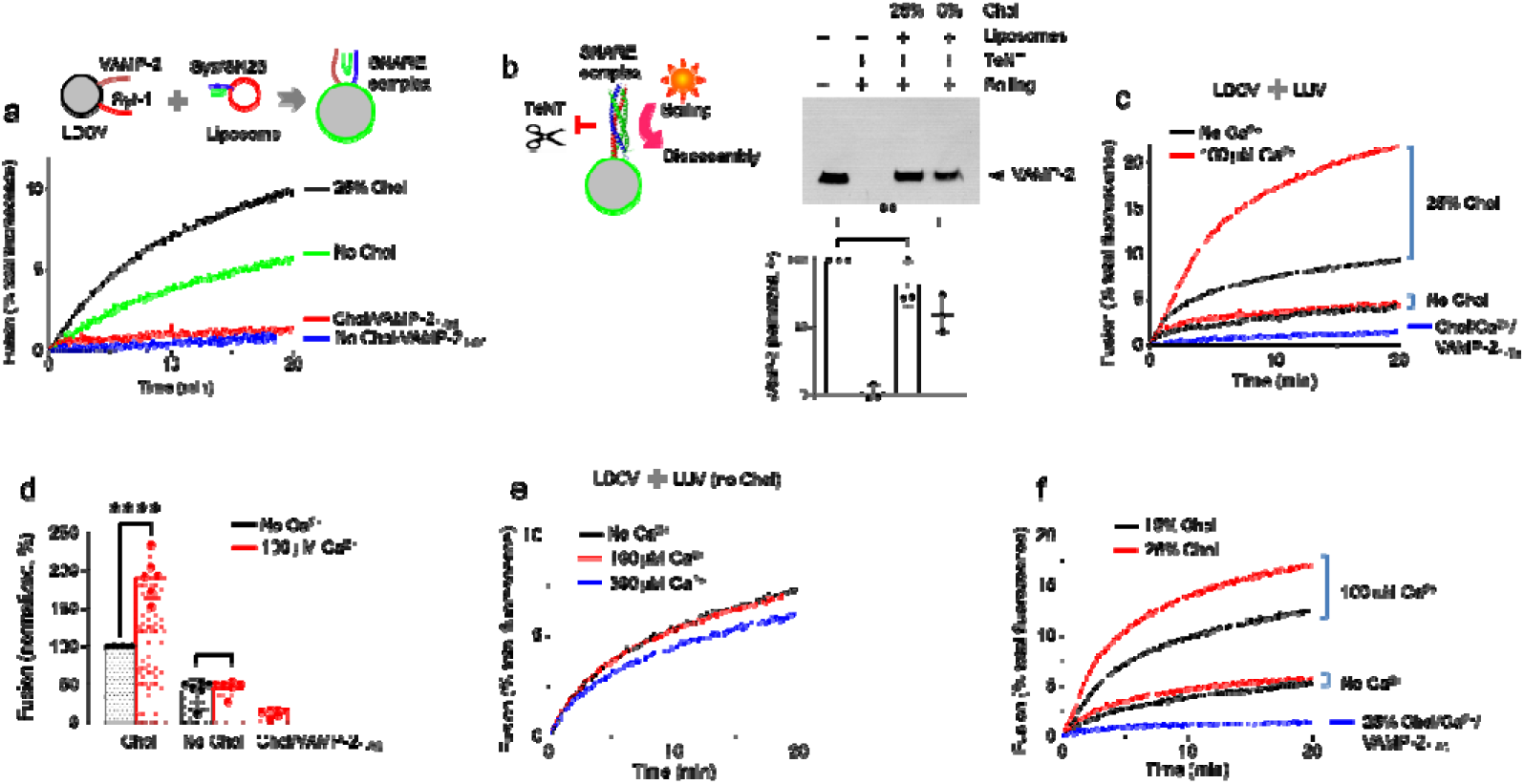
Cholesterol is required for Ca^2+^-triggered vesicle fusion. (**a**) *In-vitro* reconstitution of large dense-core vesicle (LDCV) fusion using a lipid-mixing assay. Purified LDCVs were incubated with PM-liposomes that incorporate the stabilized Q-SNARE complex of syntaxin-1A/SNAP-25A (Syx/SN25) in a 1:1 molar ratio (**Online Methods**). Cholesterol (Chol) was included either 25% or 0% in the PM-liposomes. For clarity, only endogenous VAMP-2 and synaptotagmin-1 of native LDCVs are shown and full fusion is presented. (**b**) Formation of the ternary SNARE complex after vesicle fusion in the presence or absence of cholesterol in the PM-liposomes. LDCVs were incubated with liposomes for 20 min without Ca^2+^, then treated with Tetanus neurotoxin (TeNT). TeNT-resistant VAMP-2 indicates the ternary SNARE complex formation in SDS-PAGE (**Online Methods**). Boiling at 95°C disrupts the ternary SNARE complex so that VAMP-2 migrates to its size. Data are mean ± SD from 3 independent experiments (n = 3). **, *p* < 0.01. (**c**,**d**) Dependence on cholesterol for Ca^2+^-dependent vesicle fusion. Addition of 100 µM free Ca^2+^ provoked fusion of LDCV with the PM-liposomes. Preincubation o the PM-liposomes with VAMP-2_1-96_ caused competitive inhibition that blocked SNARE-mediated fusion. Fusion is presented as a percentage of basal fusion with Chol-containing PM-liposomes in the absence of Ca^2+^. Data are mean ± SD (n = 6 independent experiments). ****, *p* < 0.0001. (**e**) Free Ca^2+^ was increased to 300 µM, when LDCVs fused with the PM-liposomes (0% Chol). (**f**) Liposomes contained either 15% or 25% of Chol. Lipid composition of the PM-liposomess: 45% PC, 15% PE, 10% PS, 25% Chol, 4% PI, and 1% PI(4,5)P_2_. When Chol was reduced, PC contents were adjusted accordingly. Physiological ionic strength and 1 mM MgCl_2_/3 mM ATP were included in all experiments.

Then we investigated the cholesterol effect on Ca^2+^-dependent fusion. Although cholesterol controls Ca^2+^-dependent neurotransmission and exocytosis in diverse cell types^8, 13, 14, 15, 16, 17, 18^, the step at which vesicle fusion is impaired has not been determined. We used native vesicles to completely reconstitute vesicle fusion that reproduces Ca^2+^-dependent vesicle fusion, correlating with the *in-vivo* data in a physiological ionic environment^31, 32, 33^. Addition of 100 µM free Ca^2+^ accelerated vesicle fusion in the presence of Mg^2+^/ATP (**Fig. 2c,d**), where synaptotagmin-1 interacts only with PIP_2_-containing membranes, but not SNARE proteins^31, 32^. VAMP-2_1-96_, the soluble cytoplasmic region of VAMP-2, blocked vesicle fusion; this result supports SNARE-dependent vesicle fusion. Surprisingly, Ca^2+^-evoked LDCV fusion was completely abolished when cholesterol in the PM-liposomes was absent (**Fig. 2c,d**). The Q-SNARE proteins were incorporated in the PM-liposomes independently of cholesterol (**Supplementary Fig. 1**). We also confirmed that Ca^2+^-dependent LDCV fusion did not occur without cholesterol (0% Chol) in the PM-liposomes when the full-length syntaxin-1A and SNAP-25A binary acceptor complex (see **Online Methods**) was included (**Supplementary Fig. 2**). High concentration of free Ca^2+^ failed to increase vesicle fusion when the PM-liposomes contained no cholesterol (**Fig. 2e**) and cholesterol accelerated Ca^2+^-dependent LDCV fusion in a dose-dependent manner (**Fig. 2f**). Altogether, our results confirm that cholesterol is required for Ca^2+^-dependent vesicle fusion and that this *in-vitro* reconstitution reproduces physiological roles of cholesterol on vesicle fusion and exocytosis.

### Cholesterol is not required for liposome-liposome fusion

We further tested SVs purified from mice brain to determine whether cholesterol is essential for Ca^2+^-dependent SV fusion. Ca^2+^ failed to accelerate SV fusion in the absence of cholesterol in the PM-liposomes (0% Chol) (**Fig. 3a-c**); this result is consistent with the cholesterol requirement for Ca^2+^- dependent LDCV fusion (**Fig. 2d**), however, Ca^2+^-independent basal SV fusion was not affected (**Fig. 3c**). A unique and major advantage of native vesicles for a fusion assay is that they maintain the native lipid and protein diversity as well as the structural integrity of vesicles to mimic endogenous vesicle fusion. Instead of purified native vesicles, vesicle-mimicking liposomes (V-liposomes) have been used for a fusion assay to study the molecular mechanisms. We therefore tested V-liposomes that incorporate full length VAMP-2 and synaptotagmin-1 to determine whether the dependence on cholesterol for Ca^2+^- dependent fusion can be reproduced (**Fig. 3d-f**). Intriguingly, Ca^2+^-dependent liposome fusion was slightly reduced but still observable even in the absence of cholesterol in the PM-liposomes (**Fig. 3d-f**). These results indicate that cholesterol is required for Ca^2+^-dependent fusion of native vesicles, i.e., LDCV and SV, but not for liposome-liposome fusion.

**Figure 3.**
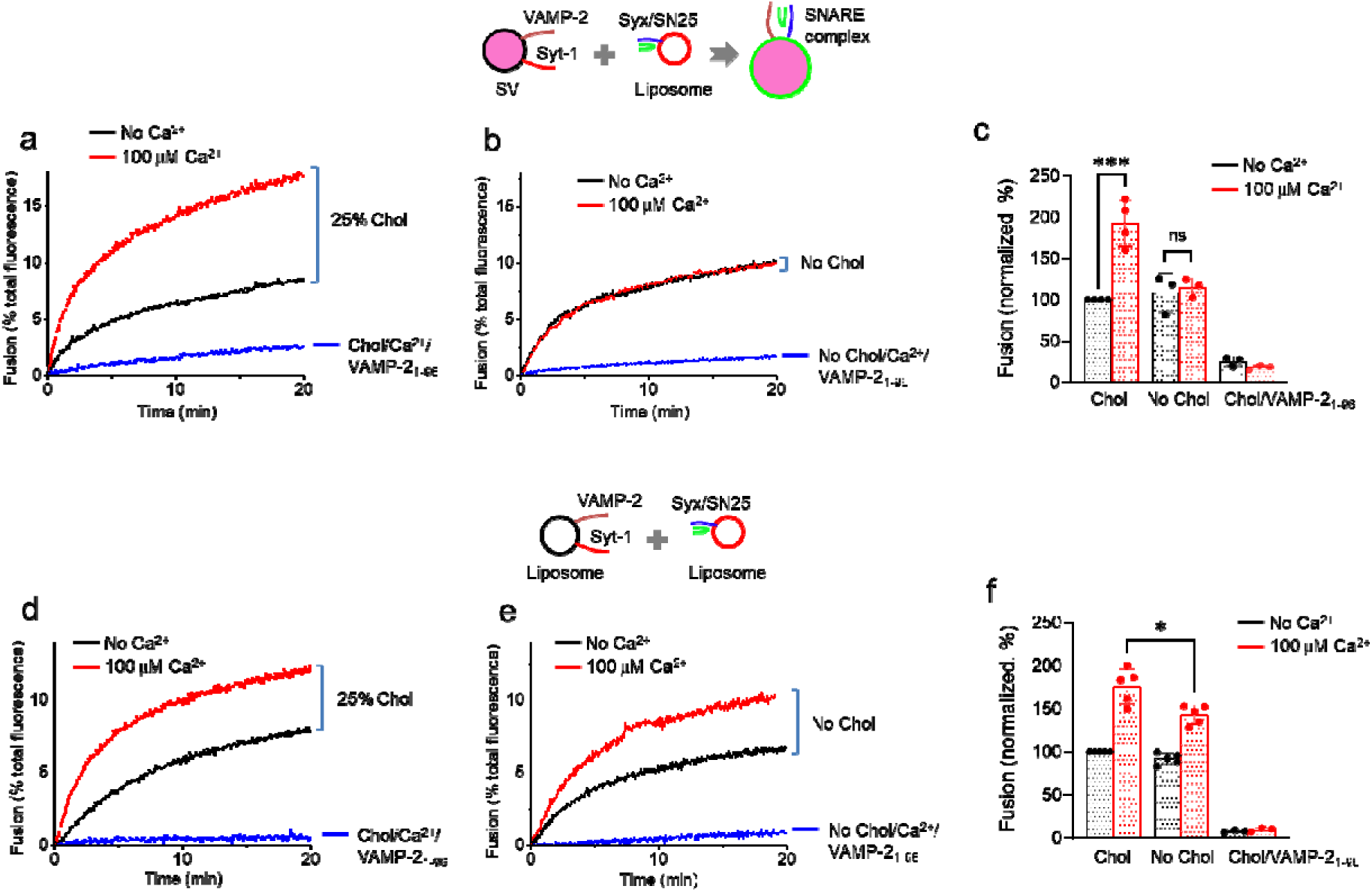
Cholesterol is not required for liposome-liposome fusion. (**a-c**) Synaptic vesicle (SV) fusion with the PM-liposomes that contain either 25% (**a**,**c**) or 0% (**b**,**c**) Chol. Only endogenous VAMP-2 and synaptotagmin-1 of native SVs are shown and full fusion is presented for clarity. (**d-f**) Instead of native LDCVs or SVs, V-liposomes that incorporated the full-length synaptotagmin-1 and VAMP-2 were incubated with PM-liposomes that contained either 25% (**d**) or 0% (**e**) Chol. Both V-liposomes and PM-liposomes are LUVs. Lipid composition of the PM-liposomes is described in **Figure 2**. Lipid composition of V-liposomes: 55% PC, 20% PE, 15% PS, and 10% Chol. (**c**,**f**) Fusion is presented as a percentage of basal fusion with Chol-containing PM-liposomes in the absence of Ca^2+^. Data in **c** and **f** are mean ± SD from 3∼5 independent experiments (n = 3∼5). *, *p* < 0.05. ***, *p* < 0.0005.

### Membrane binding of synaptotagmin-1 independently of cholesterol

Cholesterol in the PM-liposomes is indispensable for Ca^2+^-dependent vesicle fusion, so we investigated whether the binding of synaptotagmin-1 to the PIP_2_-containing membrane is affected by cholesterol. The C2AB domain of synaptotagmin-1 (Syt_97-421_) was labelled with Alexa Fluor 488 at S342C as a donor, and the PM-liposomes (Lip., protein-free) were labelled with Rhodamine (Rho)-PE as an acceptor (**Online Methods**). The C2AB binding to liposomes was monitored by FRET between the C2AB domain (Alexa Fluor 488) and Rhodamine-labelled PM-liposomes (**Fig. 4a**). The C2AB domain of synaptotagmin-1 could bind to both cholesterol-containing and cholesterol-free liposomes in response to Ca^2+^, and the Ca^2+^ titration for C2AB binding to the PM-liposomes (0% Chol) was comparable to that of 25% Chol-containing PM-liposomes; this result demonstrates Ca^2+^-dependent C2AB binding to anionic phospholipids regardless of cholesterol in liposomes (**Fig. 4a-c**). The PM-liposomes contain anionic phospholipids including 10% PS, 4% PI, and 1% PIP_2_, which provide complete coordination sites for the Ca^2+^-bound C2AB domain to interact with the membranes. (**Fig. 4a-c**).

**Figure 4.**
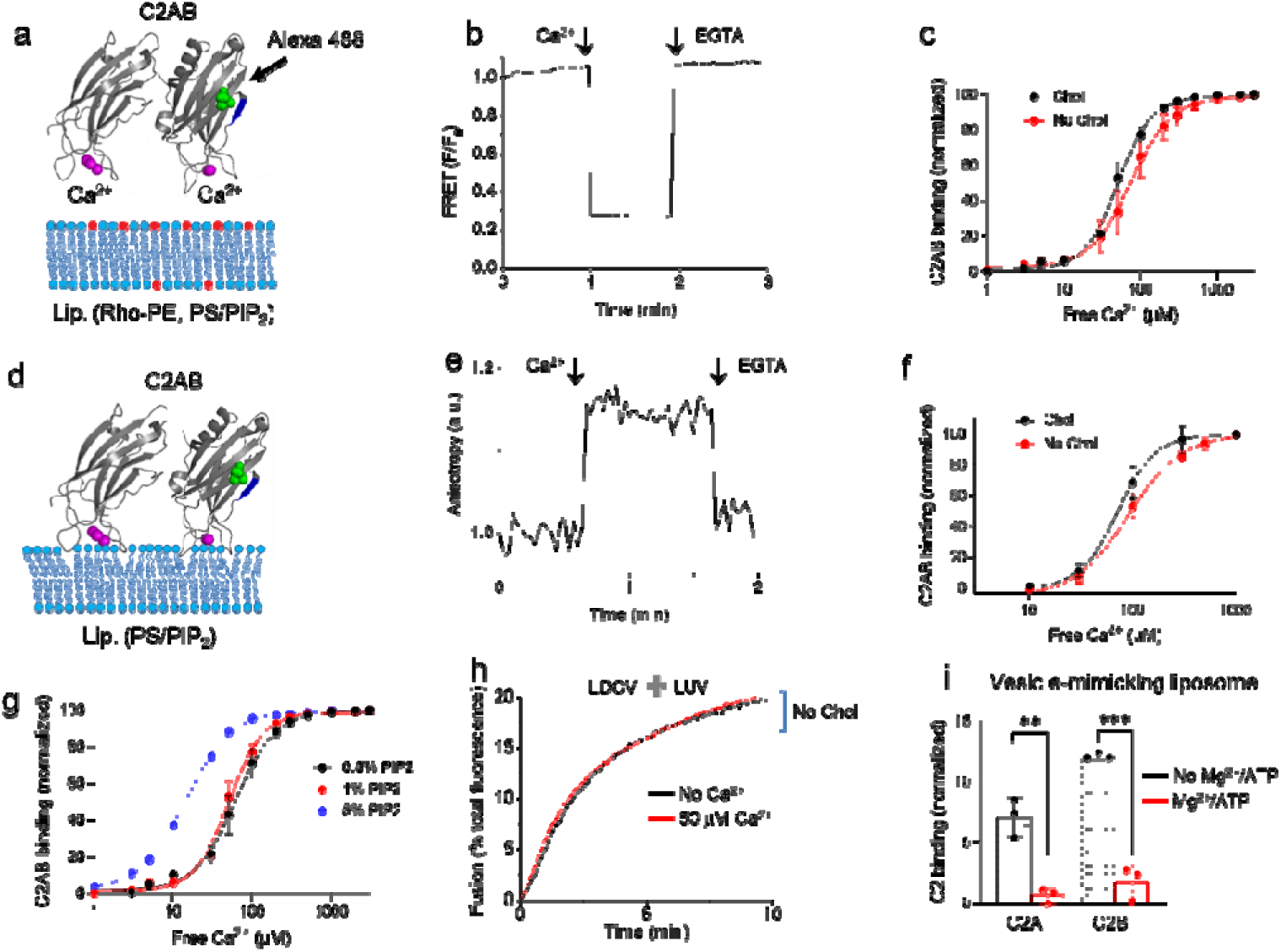
Membrane binding of the C2AB domain of synaptotagmin-1 in the presence or absence of cholesterol. (**a-c**) Membrane binding of the C2AB domain of synaptotagmin-1 was monitored using FRET in which the C2AB domain (Syt-1_97-421_) was labelled with Alexa Fluor 488 at S342C (green dots) as a donor, and the PM-liposomes (Lip.) were labelled with Rhodamine (Rho)-PE (red) as an acceptor (**Online Methods**). The PM-liposomes (protein-free) contained either 25% (**b**) or 0% Chol. (**c**) C2AB binding is presented as a percentage of maximum C2AB binding induced by 1 mM free Ca^2+^. Data are mean ± SD from 3∼5 independent experiments. (**d-f**) Binding of the C2AB domain to the PM-liposomes (protein-free; lipid composition as in **a-c** without Rho-PE) was monitored using fluorescence anisotropy. (**f**) Dose-response curve of Ca^2+^-dependent C2AB binding to the PM-liposomes that contain either 25% or 0% Chol. Data are mean ± SD (n = 3∼4 independent experiments). (**g**) FRET was conducted to monitor C2AB binding to liposomes as in **Fig. 4a-c**. PIP_2_ was incorporated in the PM-liposomes (25% cholesterol included, protein-free). (**g**) High PIP_2_ concentration increased Ca^2+^ sensitivity for C2AB binding to membranes. Data are mean ± SD (n = 3∼5 independent experiments). (**h**) LDCV fusion with the PM-liposomes that contain 5% PIP_2_ concentration and no Chol. (**i**) C2AB binding to V-liposomes was monitored using a tryptophan-dansyl FRET pair. Neither the C2A nor C2B domain binds t V-liposomes (no PIP_2_) by 1 mM Ca^2+^ in the presence of 1 mM MgCl_2_/3 mM ATP. Data are mean ± SD (n = 3 independent experiments). **, *p* < 0.01. ***, *p* < 0.0005.

To further assess membrane binding of the C2AB domain, a fluorescence anisotropy measurement was conducted to monitor the rotational mobility of the C2AB domain, labelled with Alexa Fluor 488 at 342C; the PM-liposomes are label-free and protein-free (**Fig. 4d**). The C2AB binding to the PM-liposomes results in the increase of fluorescence anisotropy due to a reduction in the rotational mobility of the membrane-bound C2AB domain (**Fig. 4e**). Ca^2+^ titration for membrane binding of the C2AB domain showed no difference between the two sets of PM-liposomes (0% vs 25% Chol) (**Fig. 4d-f**). The increases of PIP_2_ concentration in the PM-liposomes shifted Ca^2+^ titration curves to the left, which indicates an increase in the Ca^2+^-sensitivity of C2AB membrane binding (**Fig. 4g, Supplementary Fig. 3**). High PIP_2_ concentration increases Ca^2+^ sensitivity for vesicle fusion^31^; this interaction implies that increase in the magnitude of the negative electrostatic potential in the plasma membranes increases its attraction of Ca^2+^-bound synaptotagmin-1. Even 5% PIP_2_ in the PM-liposomes Ca^2+^ still failed to increase LDCV fusion without cholesterol (**Fig. 4h**).

We have previously reported the membrane binding of the C2AB domain in a physiological ionic environment^31, 32^, showing that the C2AB domain can only bind to the PIP_2_-containing plasma membrane, not to the SNARE complex. Then, we further validated if either the C2A or C2B domain still binds to vesicle membrane at physiological ionic strength. Indeed, the C2A and C2B domains bind to V-liposomes (0% PIP_2_, 15% PS included), but this interaction was completely disrupted at physiological ionic strength due to the charge-shielding effect of Mg^2+^ and ATP (**Fig. 4i**)^31, 32^. Both the C2A and C2B domains bind to the plasma membrane to drive vesicle fusion in a physiological ionic environment^31, 32^, proposing that membrane binding of both the C2A and C2B domains might have cooperative and synergistic effect on membrane deformation. Taken together, C2AB binding into the plasma membrane directly provides the driving force to lower the energy barrier for Ca^2+^-triggered vesicle fusion to occur. Cholesterol is only required for Ca^2+^-dependent vesicle fusion, but is not essential for Ca^2+^-dependent C2AB membrane binding.

### Membrane curvature is critical for Ca^2+^-dependent fusion

Despite the complete inhibition of Ca^2+^-dependent vesicle fusion (**Fig. 2 and 3a,b**), membrane binding of the C2AB domain was still observable in the absence of cholesterol in liposomes (**Fig. 4**); this observation strongly suggests that the downstream of membrane binding of synaptotagmin-1 is disrupted. Hydrophobic residues in the Ca^2+^-binding loops of synaptotagmin-1 penetrate the inner leaflet of the plasma membrane^35^, and probably lead to local membrane bending and deformation^36, 37^. This local membrane deformation might accelerate vesicle fusion by lowering the energy barrier^38^. Cholesterol also regulates local membrane bending and deformation^39^, so in the next experiments we examined whether cholesterol has a critical function in membrane bending to trigger Ca^2+^-dependent vesicle fusion.

To this end we used different PM-liposomes of different sizes: large (LUV) with 110-nm diameter and small unilamellar vesicles (SUV) with 60-nm diameter (**Online Methods, Supplementary Fig. 5**). The average diameter of LDCVs is 150 nm, ranging from 100 nm to 300 nm^33^.We expected that the synaptotagmin-1-induced changes in local membrane curvature and tension would be minimized in small liposomes (**Fig. 5a,b**), because they are already highly curved^37^. Indeed, Ca^2+^-dependent LDCV fusion was dramatically lower when SUVs were used compared to LUVs (**Fig. 5c-e**). The curvature effect on vesicle fusion was reproduced when the concentration of liposomes and the ratio of vesicle to liposome were changed (**Supplementary Fig. 4a-d**).

**Figure 5.**
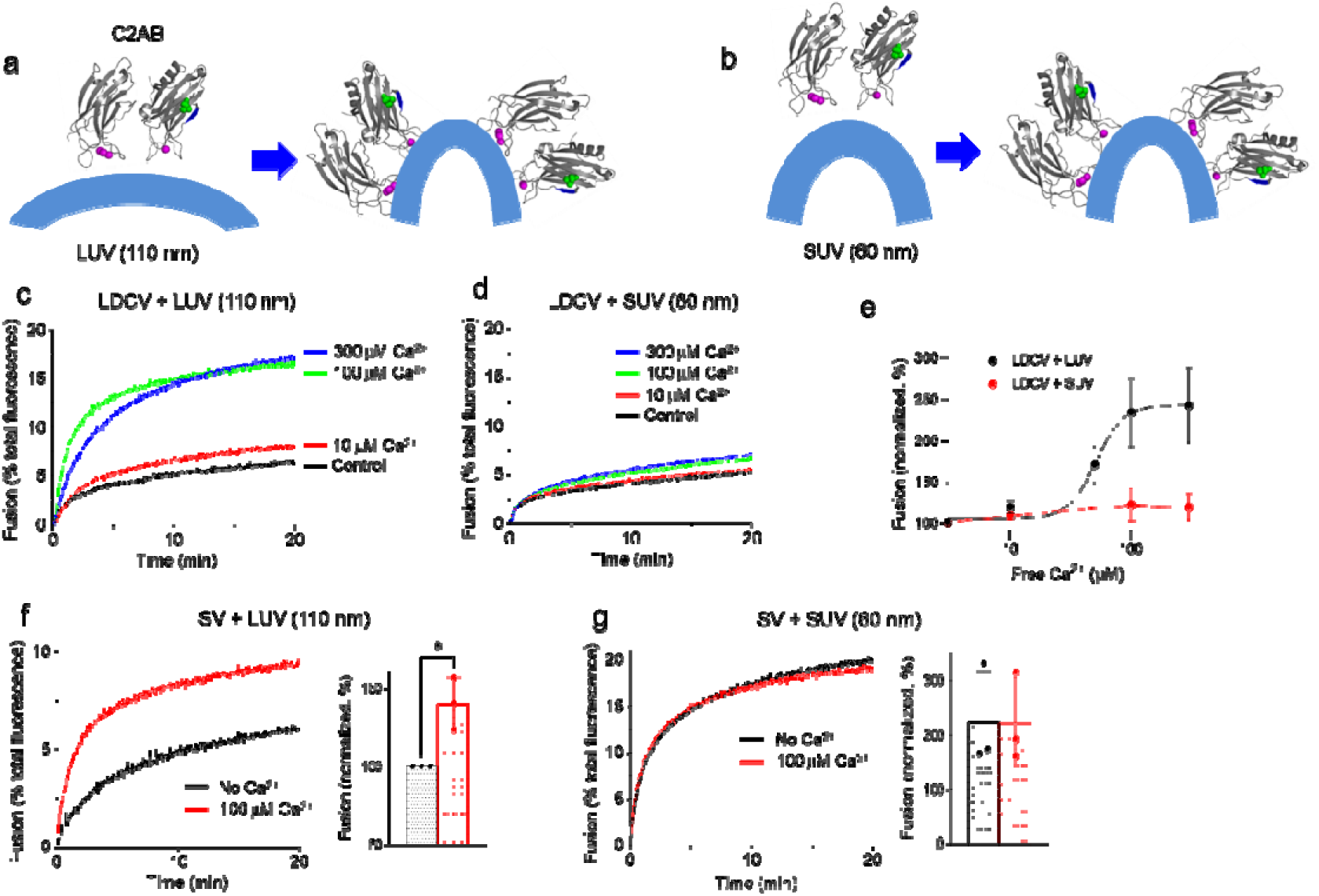
High curvature strain of membrane increases basal fusion but decreases Ca^2+^-triggered fusion. (**a**,**b**) Schematic diagram showing membrane bending by membrane binding of the C2AB domain. Large unilamellar vesicles (LUVs), 110 nm in diameter; small unilamellar vesicles (SUVs), 60 nm in diameter. The SUV is already highly curved. (**c**,**d**) Ca^2+^-dependent LDCV fusion with the PM-liposomes, either LUV or SUV. (**e**) Dose-response curve of Ca^2+^ on LDCV fusion with either LUV or SUV. (**f**,**g**) SV fusion with the PM-liposomes, either LUV or SUV. Mouse SVs are ∼45 nm in diameter. Data in **e**,**f**,**g** are mean ± SD (n = 3 ∼ 7 independent experiments). *, *p* < 0.05.

To further assess the effect of membrane curvature on vesicle fusion, we used SVs that have an average diameter of 45 nm^40^. As expected, Ca^2+^-dependent SV fusion with SUVs was completely impaired (**Fig. 5f,g**), whereas basal SV fusion was already saturated and augmented (**Fig. 5g**). Next, we replaced native SVs with small liposomes to confirm the curvature effect on vesicle fusion; small V-liposomes (SUV) that incorporate full length VAMP-2 and synaptotagmin-1 fuse with small PM-liposomes (SUV) either in the presence or absence of cholesterol (**Supplementary Fig. 4e,f**). Indeed, no Ca^2+^-dependent fusion was observed in case of small V-liposomes and small PM-liposomes, whereas LUVs show Ca^2+^- dependent liposome fusion (**Fig. 3d-f**). Altogether, high membrane tension and curvature elevate basal fusion, but decrease Ca^2+^-dependent vesicle fusion, because highly curved-membranes are likely to fuse without Ca^2+^ and show less change in membrane tension by membrane insertion of synaptotagmin-1.

### Cholesterol stabilizes membrane bending and deformation

The C2AB domain of synaptotagmin-1 has tubulation activity by membranes deformation and curvature generation^36, 37^. Next, we performed a tubulation assay using negative stain TEM to test that cholesterol regulates local membrane bending and deformation induced by synaptotagmin-1 (**Fig. 6**). As expected, Ca^2+^/C2AB domain induced tubulation by deforming membrane in the presence of cholesterol, whereas either Ca^2+^ or the C2AB domain alone had no effect (**Fig. 6a-d**), as correlating with the previous reports^36, 37^. Intriguingly, we observed no tubulation activity of the Ca^2+^/C2AB domain in the absence of cholesterol in the PM-liposomes (**Fig. 6c,d**). These TEM data provide evidence that cholesterol is crucial for stabilizing and strengthening membrane bending and deformation caused by synaptotagmin-1.

**Figure 6.**
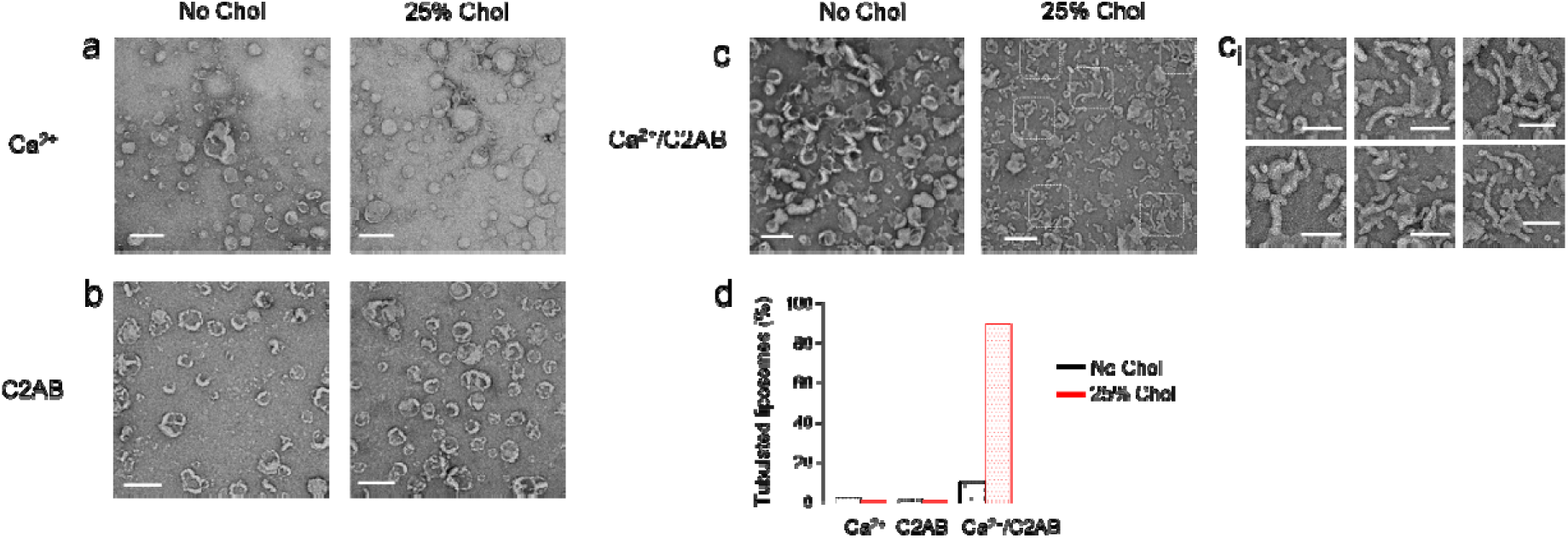
Cholesterol stabilizes and strengthens membrane deformation induced by the C2AB domain. TEM images of the PM-liposomes in the absence or presence of 25% Chol, incubated with either 100 µM Ca^2+^ (**a**), 2 µM C2AB domain (**b**), or Ca^2+^/C2AB domain (**c**). Scale, 200 nm. (**c**_**i**_) High magnification images of tubulated liposomes (white dotted lines in the right panel of **c**). Scale, 100 nm. (**d**) Quantification of tubulated liposomes (n = 110 ∼ 475 liposomes in each condition from three independent experiments).

## Discussion

Cholesterol regulates vesicle fusion as follows. First, it causes clustering of SNARE proteins in the plasma membrane. This clustering could increase the efficiency of membrane fusion^29, 30^. Cholesterol in the PM-liposomes slightly increased the formation of SNARE complexes, and thus increased LDCV fusion (**Fig. 2a-d**). Second, cholesterol regulates the physical structure, fluidity, tension, and thickness of lipid membranes^25^. Cholesterol causes negative membrane curvature that might facilitate and promote vesicle fusion^13, 25, 41, 42^. Cholesterol also contributes to vesicle fusion by stabilizing fusion pores^25, 39, 43, 44^. The plasma membrane deformations occur prior to vesicle fusion, and pre-fusion membrane curvature changes can be observed^45^. The high membrane bending energy can be released for fusion pore formation^46^. For examples, smaller vesicles, e.g., SUVs (Supplementary Fig. 4e,f) and SVs (Fig. 5f,g), have greater bending energy per unit of surface area and accordingly show robust basal fusion efficiency compared to large vesicles. Here, our data provide a novel model in which cholesterol mediates Ca^2+^-dependent vesicle fusion by stabilizing local membrane bending caused by synaptotagmin-1 binding. The curved plasma membrane has high bending energy, which can be released to drive fusion with vesicle membranes^36, 37, 46^. Thus, high membrane-bending energy stabilized by cholesterol lowers the energy barrier for vesicle fusion, thereby triggering Ca^2+^-dependent vesicle fusion (**Fig. 7a,b**).

**Figure 7.**
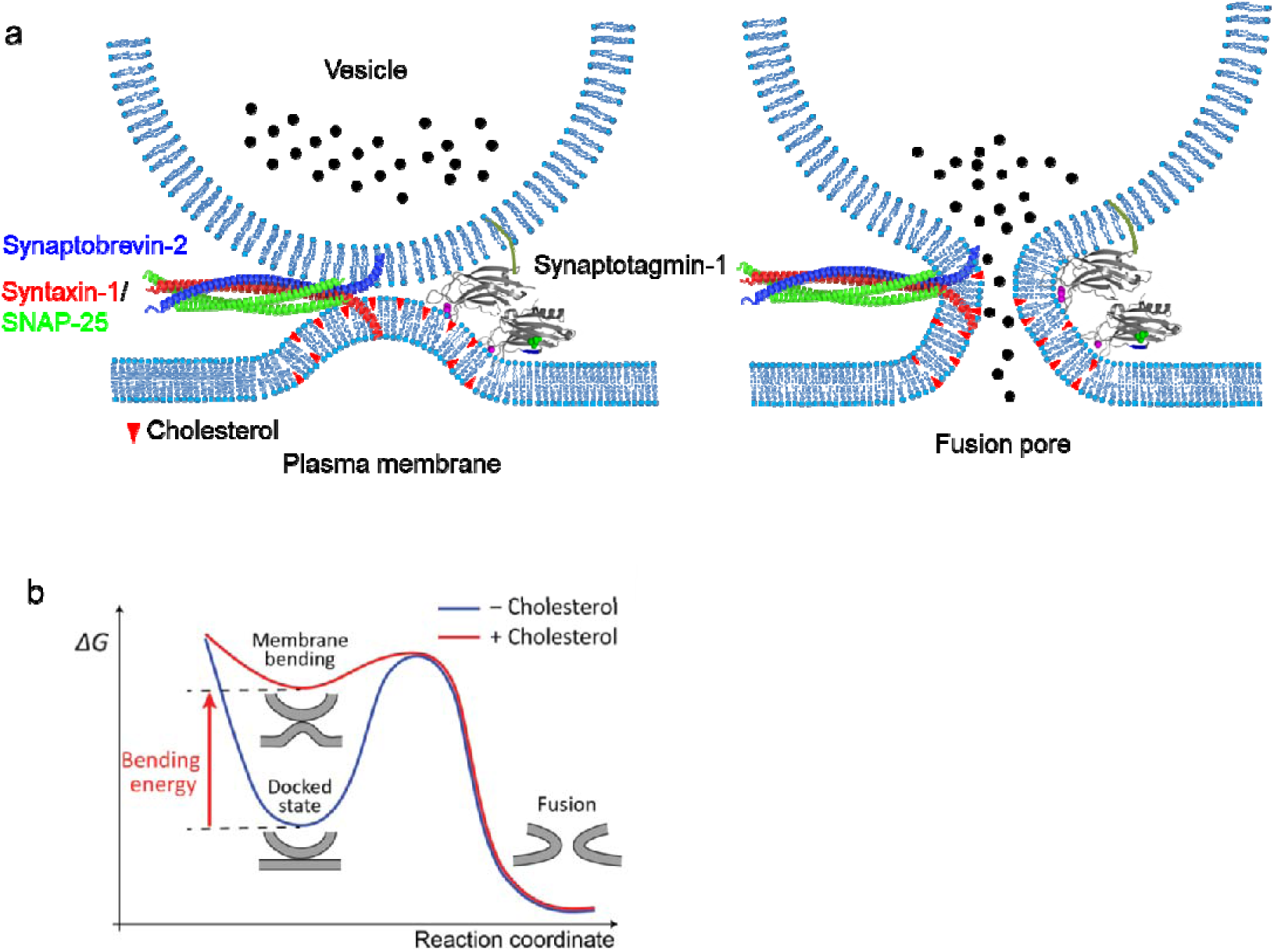
A schematic diagram summarizing synaptotagmin-1-induced membrane bending for Ca^2+^-dependent vesicle fusion. (**a**) Cholesterol stabilizes the plasma membrane deformation and bending caused by synaptotagmin-1. (**b**) Energy landscapes of vesicle fusion by synaptotagm n-1. Synaptotagmin-1-induced membrane bending enhanced by cholesterol generates the significant membrane bending energy that can drive the membrane fusion by lowering the energy barriers to promote the fusion intermediate.

The C2AB domain of synaptotagmin-1 is inserted into the plasma membrane^35^ and the membrane-binding energy of the C2AB domain is ∼18 k_B_T;^47^ the binding leads to local membrane bending and deformation^36, 37^. Membrane bending by synaptotagmin-1 could accelerate fusion by reducing the energy barrier. However, our data indicate that membrane bending by synaptotagmin-1 seems not enough to overcome the threshold of the energy barrier for fusion, when cholesterol is absent in the plasma membrane (**Fig. 2,3, and 6**). Cholesterol stabilizes and strengthens the plasma membrane deformation and bending caused by synaptotagmin-1 (**Fig. 6d**). This significant membrane bending energy can drive the membrane fusion (**Fig. 7a,b**).

Recently, Zuber’s team observed membrane bending upon calcium influx in synaptosomes^48^. Using time-resolved cryo-electron tomography within 7 to 35 ms after diffusion of 52 mM KCl in synaptosomes, Zuber’s group reported that fusion initiation occurs by membrane curvature (‘buckling’) of synaptic vesicle and the plasma membrane, showing Ca^2+^-dependent membrane bending before full fusion^48^. This cryo-electron tomography correlate with and strongly support our model that the Ca^2+^- bound C2AB domain of synaptotagmin-1 induce membrane bending and curvature for vesicle fusion.

Our *in-vitro* reconstitution of vesicle fusion has an advantage of using purified native vesicles, i.e., LDCVs and SVs, and reproduces the physiological cholesterol effect on vesicle fusion, whereas V-liposomes fail (**Fig. 3d-f**). V-liposomes are independent of cholesterol for Ca^2+^-dependent fusion, in contrast to native vesicles (**Fig. 2**,**3**). V-liposomes might have some limitations to fully replace native vesicles for a fusion assay. Native vesicles differ from V-liposomes in lipid composition, protein density, contents diversity, and physical property; therefore the structural integrity of native vesicles allows them to mimic endogenous vesicle fusion in the *in-vitro* reconstitution. For example, we observed that native vesicles, LDCVs and SVs, remain stable and functionally active even after 3∼5 times freeze-thaw cycles^31, 33^. Despite 3∼5 times snap freeze-thaw cycles, biophysical and biochemical properties of purified LDCVs are preserved; i) the structure and size distribution of LDCVs are normal, ii) complete SNARE-dependent fusion of LDCVs using a lipid-mixing assay, and iii) acidification of LDCVs is fully functional^33^, suggesting that the membrane stability and rigidity of native vesicles after freeze-thaw cycles remain stable and functional. In contrast, proteoliposomes become immediately disrupted and destructed by freezing regardless of cholesterol in liposomes, thus these liposomes fail to undergo fusion (data not shown). Liposomes should not be frozen, because the freezing process fractures or ruptures liposomes, leading to destruction of liposome membranes; i.e., ice crystals generated by the freeze-thaw process cause liposome membranes to rupture. Ice crystal formation in the outer parts of liposomes can cause the interior of liposomes to expand until liposome membrane bursts.

However, vesicular proteins such as scaffold proteins and granins could stabilize vesicle structure and regulate membrane rigidity^49^. Native vesicles have high membrane rigidity compared to V-liposomes and native vesicles seem to require higher membrane bending energy than V-liposomes do, in order to overcome the barrier for fusion due to membrane fluidity and rigidity. The structural integrity of native vesicles can be advantages for the *in-vitro* reconstitution of vesicle fusion to reproduce physiological exocytosis, but the cause of the difference between native vesicles and liposomes for fusion efficiency, and the reproduction of physiological vesicle fusion remain topics for further study.

The molecular mechanisms of synaptotagmin-1 to trigger Ca^2+^-dependent fusion remain controversial; at least six different competing models have been proposed^21^. Synaptotagmin-1 mediates Ca^2+^- dependent fusion by the electrostatic interaction, so several different synaptotagmin-1 models have been proposed, depending on the ionic environment^32^. Here we confirmed the cholesterol effect on vesicle fusion in a physiological ionic environment, i.e., normal ionic strength with Mg^2+^/ATP. Both the C2A and C2B domains of synaptotagmin-1 are inserted into the plasma membrane (**Fig. 4i**) without interacting with SNARE proteins^32^, and therefore lead to membrane bending and deformation (**Fig. 6**). Ca^2+^ fails to trigger fusion in the absence of cholesterol despite the proteins, both SNARE assembly and membrane binding of synaptotagmin-1, being fully active and functional (**Fig. 2, 4**). Cholesterol has an critical function in Ca^2+^-dependent fusion, as an important lipid regulator by stabilizing membrane deformation and curvature caused by synaptotagmin-1.

AgeL□Jrelated cholesterol reduction is linked to reduced synaptic activity, and defects in synaptic transmission by cholesterol deficiency could result in neurodegeneration^4^. Our data explains the molecular mechanisms how cholesterol contributes to synaptic transmission and neuronal function and may pave the way for development of studies to explore to treatment of neurodegenerative and neurodevelopmental disorders by optimizing cholesterol levels in the plasma membrane.

## Acknowledgements

We thank Dr. Reinhard Jahn for constructs and samples. We are deeply indebted to Dr. Kyong-Tai Kim for technical assistanace. We thank Drs. Yongfeng Tong and Akshath Raghu Shetty from the Core Labs of the Qatar Environment and Energy Research Institute (QEERI) for technical support. Thanks to Dr. Ahmed Elalawy and Sarra Karrar from Widam Food Company for the arrangement of adrenal glands. This work was supported by the grant from Qatar Biomedical Research Institute (Project Number SF 2019 004 to Y.P.).

## Figure legends

**Supplementary Figure 1.**
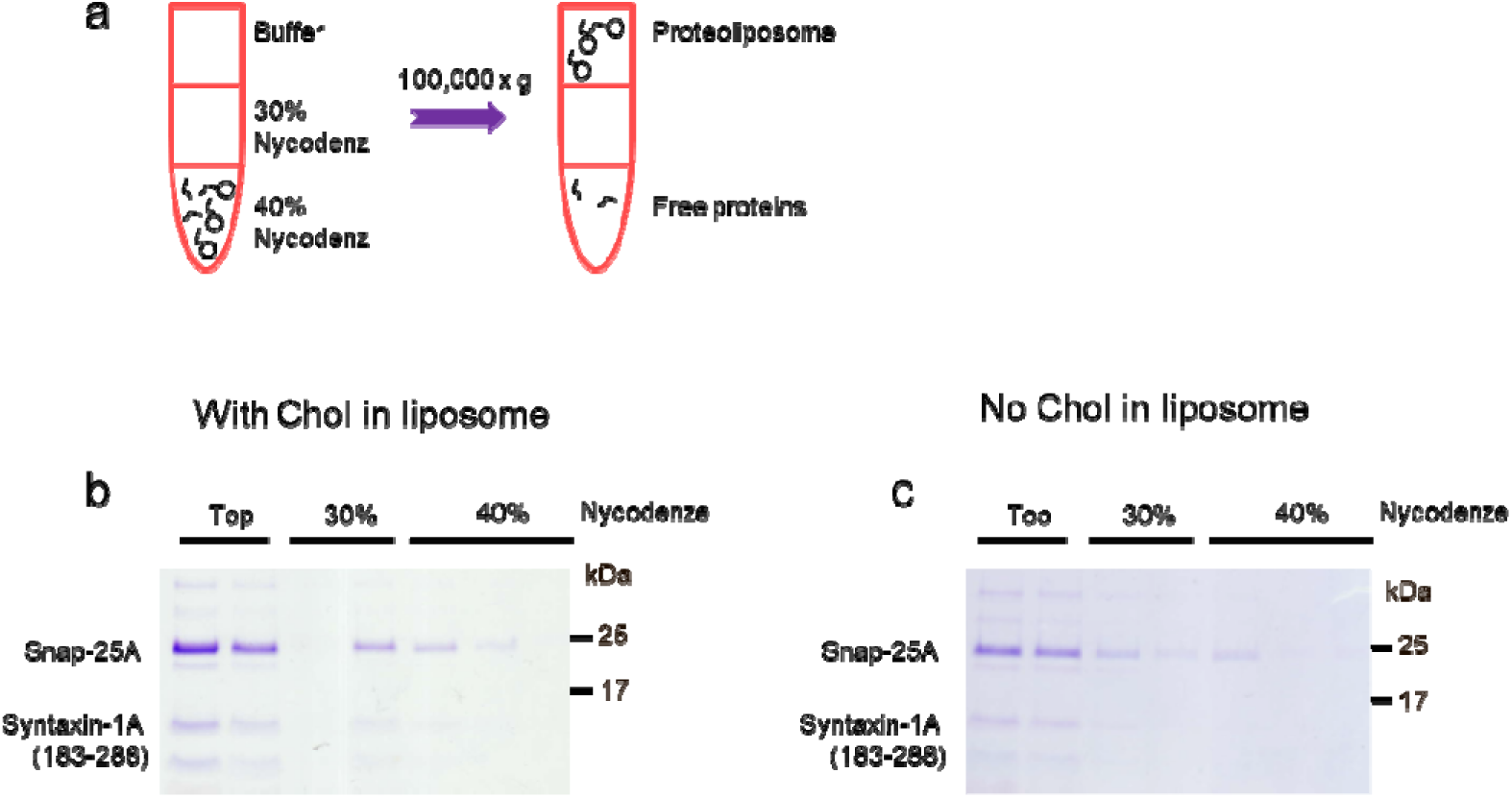
Incorporation of the Q-SNARE complex in liposomes. (**a**,**b**) Incorporation of the Q-SNARE complex in liposomes in the presence or absence of cholesterol was tested using a flotation assay. (**a**) Schematic diagram showing that liposomes float up through the gradient due to their buoyancy, whereas free proteins remain in the bottom of the gradient. (**b**) SNARE proteins stained by coomassie blue dyes. The Q-SNARE proteins SNAP-25A (no cysteine, cysteines are replaced by alanines) and syntaxin-1A (aa 183–288) in a 1:1 ratio by the C-terminal VAMP-2 fragment (aa 49–96), are incorporated in liposomes independently of cholesterol.

**Supplementary Figure 2.**
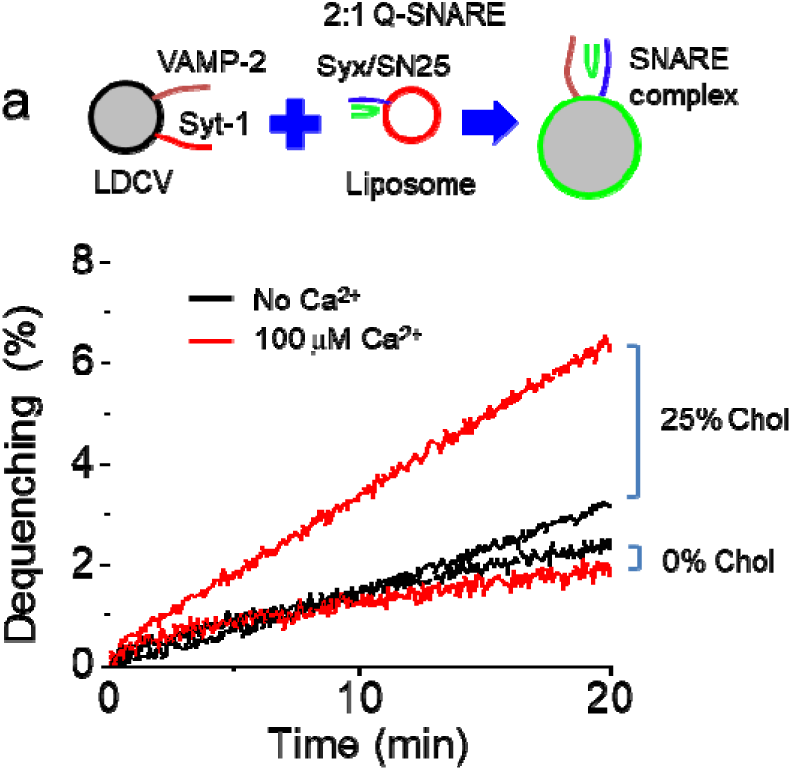
Cholesterol requirement for Ca^2+^-evoked LDCV fusion. (**a**) No Ca^2+^-dependent LDCV fusion was observed when the Q-SNARE complex consisting of the full-length syntaxin-1A (1-288) and SNAP-25A (no cysteine, cysteines are replaced by alanines) was included in liposomes with (25% Chol) and without cholesterol (0% Chol). The binary Q-SNARE complex shows slow fusion rate and relatively low fusion activity, as expected^31, 34^.

**Supplementary Figure 3.**
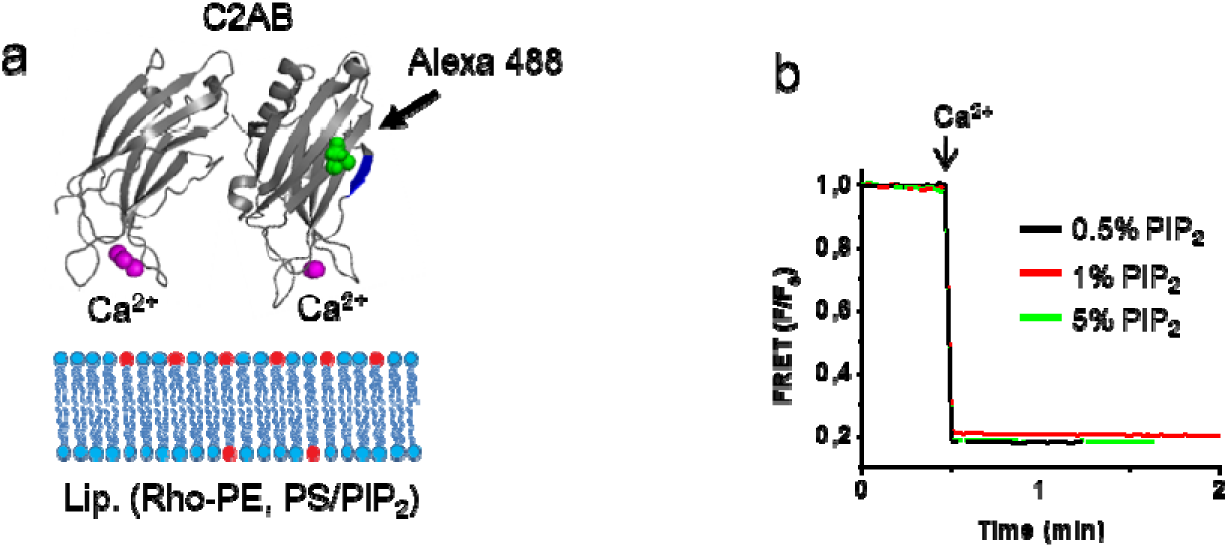
Monitoring C2AB binding to liposomes using FRET measurement. (**a**) A schematic diagram of C2AB domain binding to liposomes. Membrane binding of the C2AB domain of synaptotagmin-1 was monitored using FRET in the C2AB domain (Syt-1_97-421_) was labelled with Alexa Fluor 488 at S342C (green dots) as a donor, and the PM-liposomes (Lip.) were labelled with Rhodamine (Rho)-PE (red) as an acceptor (**Online Methods**). Lipid composition of the PM-liposomes: 45% PC, 13.5% PE, 10% PS, 25% Chol, 4% PI, 1% PI(4,5)P_2_, and 1.5% Rho-PE. When PI(4,5)P_2_ was changed, PI contents were adjusted accordingly. (**b**) PI(4,5)P_2_ contents were increased to 5% in liposomes and C2AB binding was saturated with 0.5% PI(4,5)P_2_ in liposomes. Maximum C2AB binding induced by 1 mM free Ca^2+^ in the presence of 1 mM MgCl_2_/3 mM ATP.

**Supplementary Figure 4.**
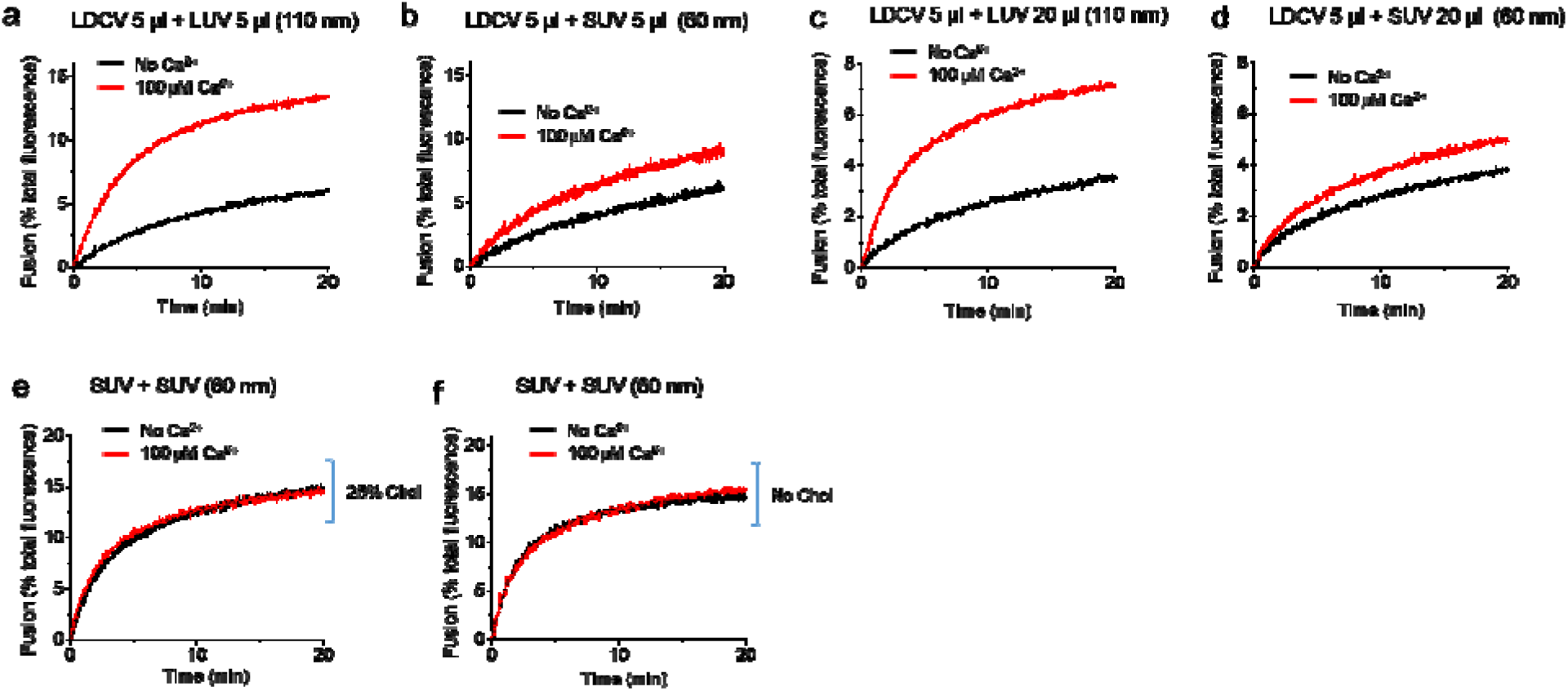
Ca^2+^-dependent vesicle fusion depends on liposome membrane curvature. Large unilamellar vesicles (LUVs), 110 nm in diameter; small unilamellar vesicles (SUVs), 60 nm in diameter. SUV is already highly curved. A lipid-mixing assay to monitor Ca^2+^-dependent LDCV fusion with the PM-liposomes, either LUV or SUV. Equal amount of LDCVs (5 µL) was incubated to induce fusion with either 5 µL (**a**,**b**) or 20 µL (**c**,**d**) LUV or SUV in 1 mL fusion buffer; 120 mM K-glutamate, 20 mM K-acetate, 20 mM HEPES-KOH (pH 7.4), 1 mM MgCl_2_, and 3 mM ATP. 100 µM free Ca^2+^ in presence of 1 mM MgCl_2_/3 mM ATP. (**e**,**f**) Both V-liposomes and PM-liposomes are SUVs the V-liposomes that incorporated the full-length synaptotagmin-1 and VAMP-2 were incubated with PM-liposomes that contained either 25% (**e**) or 0% (**f**) Chol. Lipid composition of the PM-liposomes and V-liposomes is described in **Figure 2** and **Figure 3**, respectively.

**Supplementary Figure 5.**
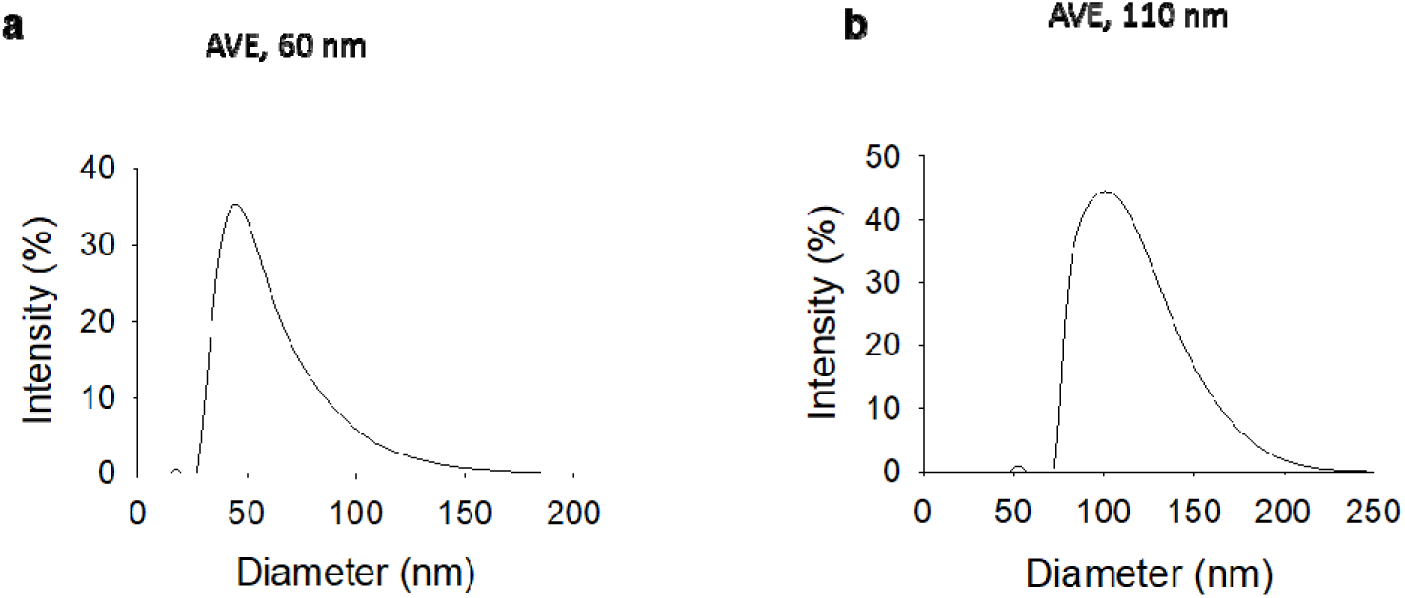
The size distribution of proteoliposomes. The size distribution of proteoliposomes was determined using dynamic light scattering (DLS). LUVs (110 nm in diameter) and SUVs (60 nm in diameter) were prepared by extrusion (**Online Methods**).

## Online Methods

### Purification of large dense-core vesicles (LDCVs) and synaptic vesicles (SVs)

LDCVs, also known as chromaffin granules, were purified from bovine adrenal medullae using continuous sucrose gradient and resuspended with fusion buffer containing 120 mM K-glutamate, 20 mM K-acetate, and 20 mM HEPES.KOH, pH 7.4, as elsewhere^33^. SV from mouse brains were purified as described elsewhere^50^. Briefly, mice brains were homogenized in homogenization buffer supplemented with protease inhibitors, using a glass-Teflon homogenizer. The homogenate was centrifuged for 10 min at 1,000g and the resulting supernatant was further centrifuged for 15 min at 15,000g. The synaptosome pellet was lysed by adding ice-cold water, followed by centrifugation for 25 min at 48,000g. The resulting supernatant was overlaid onto a 0.7 M sucrose cushion and centrifuged for 1 h at 133,000g. The pellet was resuspended in fusion buffer (120 mM K-glutamate, 20 mM K-acetate, 20 mM HEPES.KOH, pH 7.4).

### Protein purification

All SNARE and the C2AB domain of synaptotagmin-1 constructs based on rat sequences were expressed in *E. coli* strain BL21 (DE3) and purified by Ni^2+^-NTA affinity chromatography followed by ion-exchange chromatography as described elsewhere^31, 32^. The stabilized Q-SNARE complex consisting syntaxin-1A (aa 183–288), SNAP-25A (no cysteine, cysteines replaced by alanines) in a 1:1 ratio by the C-terminal VAMP-2 fragment (aa 49–96) was purified as described earlier^34^. The binary Q-SNARE complex containing the full-length syntaxin-1A (1-288) and SNAP-25A (no cysteine, cysteines replaced by alanines) was expressed using co-transformation^31^. The full-length VAMP-2, soluble cytoplasmic region of VAMP-2 (VAMP-2_1-96_), full-length synaptotagmin-1, C2AB domain of synaptotagmin-1 (aa 97-421), C2A domain (aa 96-262), C2B domain (aa 248-421), and C2ab mutant (D178A, D230A, D232A, D309A, D363A, D365A) were purified by Mono S column (GE Healthcare, Piscataway, NJ) as described previously^51^. The stabilized Q-SNARE complex and the syntaxin-1A/SNAP-25A binary SNARE complex were purified by Ni^2+^-NTA affinity chromatography followed by ion-exchange chromatography on a Mono Q column (GE Healthcare, Piscataway, NJ) in the presence of 50 mM n-octyl-β-D-glucoside (OG)^31^. The point mutated C2AB domain (S342C) was labelled with Alexa Fluor 488 C5 maleimide (C2AB^A488^)^51^.

### Lipid composition of liposomes

Lipid composition (molar percentages) of the PM-liposomes that contain the Q-SNARE complex consists of 45% PC (L-α-phosphatidylcholine, Cat. 840055), 15% PE (L-α-phosphatidylethanolamine, Cat. 840026), 10% PS (L-α-phosphatidylserine, Cat. 840032), 25% Chol (cholesterol, Cat. 700000), 4% PI (L-α-phosphatidylinositol, Cat. 840042), and 1% PI(4,5)P_2_ (Cat. 840046). When cholesterol was excluded (0% Chol), PC contents were accordingly adjusted. In case of changing PI(4,5)P_2_ concentration, PI contents were accordingly adjusted. VAMP-2/synaptotagmin-1-containing V-liposomes are composed of 55% PC, 20% PE, 15% PS, and 10% Chol. For FRET-based lipid-mixing assays, 1.5% 1,2-dioleoyl-sn-glycero-3-phosphoethanolamine-N-(7-nitrobenz-2-oxa-1,3-diazol-4-yl (NBD-DOPE) and 1.5% 1,2-dioleoyl-sn-glycero-3-phosphoethanolamine-N-lissamine rhodamine B sulfonyl ammonium salt (Rhodamine-DOPE) were incorporated in the PM-liposomes (accordingly 12% unlabeled PE) as a donor and an acceptor dye, respectively. For FRET measurement using C2AB^A488^, 1.5% Rhodamine-DOPE was included in the PM-liposomes (protein-free). In the case of FRET for tryptophan, 5% N-(5-dimethylaminonaphthalene-1-dulfonyl)-1,2-dihexadecanoyl-sn-glycero-3-phosphoethanolamine, triethylammonium salt (dansyl-DHPE) was incorporated in the PM-liposomes (protein-free). All lipids were from Avanti Polar lipids except dansyl-DHPE (Invitrogen).

### Preparation of proteoliposomes

Incorporation of the stabilized and binary Q-SNARE complex into large unilamellar vesicles (LUVs, 110 nm in diameter) was achieved by OG-mediated reconstitution, called the direct method, i.e. incorporation of proteins into preformed liposomes^31, 32^. Briefly, lipids dissolved in a 2:1 chloroform-methanol solvent were mixed according to lipid composition. The solvent was removed using a rotary evaporator (which generated lipid film on a glass flask), then lipids were resuspended in 1.5 mL diethyl ether and 0.5 mL buffer containing 150 mM KCl and 20 mM HEPES/KOH pH 7.4. After sonication on ice (3 × 45 s), multilamellar vesicles were prepared by reverse-phase evaporation using a rotary evaporator as diethyl ether was removed. Multilamellar vesicles (0.5 mL) were then extruded using polycarbonate membranes of pore size 100 nm (Avanti Polar lipids) to give uniformly-distributed LUVs with the average diameter of 110 nm (**Supplementary Fig. 5**). After the preformed LUVs had been prepared, SNARE proteins or the full-length VAMP-2/synaptotagmin-1 were incorporated into them using OG, a mild non-ionic detergent, then the OG was removed by dialysis overnight in 1 L buffer containing 150 mM KCl and 20 mM HEPES/KOH pH 7.4 together with 2 g SM-2 adsorbent beads.

To make small unilamellar vesicles (SUVs) using the direct method, 110-nm LUVs produced as described above extruded through polycarbonate membranes with 50-nm pore size (yielding SUV that had average diameter of 60 nm, **Supplementary Fig. 5**). After preparing preformed SUVs, protein incorporation was completed by OG as described for LUV. The size distribution of proteoliposomes was determined using dynamic light scattering (DLS) (**Supplementary Fig. 5**). Proteoliposomes have protein-to-lipid molar ratio of 1:500 (n/n).

### Vesicle fusion assay

A FRET-based lipid-mixing assay was applied to monitor vesicle fusion *in vitro*^31, 32^. LDCV or SV fusion reactions were performed at 37°C in 1 mL fusion buffer containing 120 mM K-glutamate, 20 mM K-acetate, 20 mM HEPES-KOH (pH 7.4), 1 mM MgCl_2_, and 3 mM ATP. ATP should be made freshly before experiments, because ATP is easily destroyed by freezing and thawing. Free Ca^2+^ concentration in the presence of ATP and Mg^2+^ was calibrated using Maxchelator simulation program. The fluorescence dequenching signal was measured using Fluoromax (Horiba Jobin Yvon) with wavelengths of 460 nm for excitation and 538 nm for emission. Fluorescence values were normalized as a percentage of maximum donor fluorescence (total fluorescence) after addition of 0.1% Triton X-100 at the end of experiments.

### Fluorescence resonance energy transfer (FRET)

The C2AB domain of synaptotagmin-1 (30 nM, S342C) was labeled with Alexa Fluor 488, a donor dye. C2AB fragment was engineered to contain a single Cys residue (S342C) and labelled with Alexa Fluor 488. 1.5% Rhodamine-DOPE (Rho-PE), incorporated in the PM-liposomes (protein-free), was used as an acceptor. Unless otherwise stated, liposomes were LUVs prepared by the direct method. Donor fluorescence signal was measured at 37°C using Fluoromax (Horiba Jobin Yvon) with wavelengths of 488 nm for excitation and 516 nm for emission in 1 mL fusion buffer containing 120 mM K-glutamate, 20 mM K-acetate, 20 mM HEPES-KOH (pH 7.4), 1 mM MgCl_2_, and 3 mM ATP. FRET was normalized as net changes of donor fluorescence intensity and C2AB binding was presented as percentage of maximum C2AB binding induced by 1 mM Ca^2+^. In **Supplementary Figure 3**, FRET was normalized as F/F_0_, where F_0_ represents the initial value of the donor fluorescence intensity.

Liposome binding of the C2AB domain was also monitored using the tryptophan-dansyl FRET pair as a donor-acceptor dye in which dansyl-DHPE incorporated in liposomes leads to quenching of fluorescence emitted from tryptophan of the C2AB domain^52^. Then1 μM C2AB, 3 μM C2A, or 3 μM C2B was incubated with V-liposomes (protein-free). Donor fluorescence signal was measured at 37°C using Fluoromax (Horiba Jobin Yvon) with wavelengths of 295 nm for excitation and 350 nm for emission in 1 mL fusion buffer containing 120 mM K-glutamate, 20 mM K-acetate, 20 mM HEPES-KOH (pH 7.4), 1 mM MgCl_2_, and 3 mM ATP. FRET monitoring C2AB binding was normalized as a percentage of (F_0_-F)/F_0_.

### Fluorescence anisotropy measurements

The C2AB fragment (20 nM, S342C) were labelled with Alexa Fluor 488^51^. Anisotropy was measured in a Fluorolog (Horiba Jobin Yvon) at 37°C in 1 ml of buffer containing 120 mM K-glutamate, 20 mM K-acetate, and 20 mM HEPES-KOH (pH 7.4), 1 mM MgCl_2_, and 3 mM ATP. Excitation wavelength was 495 nm, and emission was measured at 520 nm. Lipid composition of the PM-liposomes (protein-free) was identical to those used in a fusion assay except labelled PE (45% PC, 15% PE, 10% PS, 25% Chol, 4% PI, and 1% PIP2). In the case of 0% Chol, PC contents were adjusted accordingly (70% PC).

### Ternary SNARE complex formation assay

Tetanus neurotoxin (TeNT) degrades free VAMP-2 whereas VAMP-2, assembled in the ternary SNARE complex, is resistant to TeNT^31^. After incubation of LDCVs with the PM-liposomes (0% or 25% Chol) that contain the stabilized Q-SNARE complex for 20 min at 37°C without Ca^2+^, the sample was subjected to TeNT treatment (200 nM, 30 min, 37°C) then boiled for 5 min at 95°C and analysed by immunoblotting with antibody against VAMP-2 (clone number 69.1, Synaptic Systems (Göttingen, Germany)).

### Preparation of bovine chromaffin cells

Chromaffin cells were isolated from the bovine adrenal gland medulla by two-step collagenase digestion as previously described^53^. The cells were grown on poly-_D_- lysine-coated glass coverslips in Dulbecco’s modified Eagle medium/F-12 (Invitrogen, CA) containing 10% fetal bovine serum (Hyclone Laboratories, UT) and 1% antibiotics (Invitrogen, CA).

### Amperometric measurement

Recordings of LDCV exocytosis from chromaffin cells were performed as described previously^53^. Carbon-fiber electrodes were fabricated from 8 μm diameter carbon fibers and back-filled with 3 M KCl. The amperometric current, generated by oxidation of catecholamine, was measured using an axopatch 200B amplifier (Axon Instruments Inc., CA), which was operated in voltage-clamp mode at a holding potential of + 650 mV. Amperometric signals were low-pass filtered at 1 kHz and sampled at 500 Hz. For data acquisition and analysis, pCLAMP 11 software (Axon Instruments) was used. The area of amperometric current represents the total amount of released catecholamine. Amperometric current generated by repetitive stimulation was individually integrated. Relative exocytosis is presented as a percentage of the first DMPP-induced total catecholamine release.

### Transmission electron microscopy (TEM)

Chromaffin cells, grown on Vitrogen collagen matrix (Cohesion, Palo Alto, CA), were washed out with Locke’s solution containing 157.4 mM NaCl, 5.6 mM KCl, 2.2 mM CaCl_2_, 1.2 mM MgCl_2_, 5.6 mM D-glucose, 5 mM HEPES, and 3.6 mM NaHCO_3,_ pH 7.4 titrated by NaOH. As described previously^53^, cells were fixed with 2% paraformaldehyde and 2% glutaraldehyde in 0.05 M sodium cacodylate buffer at pH 7.4 for 20 min at room temperature. Cells were post-fixed with 0.5% osmium tetroxide in 0.05 M sodium cacodylate buffer at pH 7.4 for 30 min at room temperature. Cells were further dehydrated in graded ethanol solutions and embedded in LR White resin (London Resin Co., Berkshire, UK). Silver-gold thin sections were stained with uranyl acetate and lead citrate. The thin sections were examined under JEOL 1200 EX2 transmission electron microscope at 80 kV.

For liposomes, 5 µL of samples were deposited on carbon-coated 400-mesh copper grids (CF400-CU, Electron Microscopy Sciences). Grids were stained with uranyl acetate for negative staining and embedded in methylcellulose-uranyl acetate. Liposomes were visualized at 80 kV in Talos F200C Transmission Electron Microscope (Thermo Fisher Scientific). The images were acquired using bottom-mounted CETA camera.

### Liposome co-flotation assay

Liposomes float up through the gradient due to their buoyancy and free proteins remain in the bottom of the gradient, whereas proteins incorporated in liposomes co-float to the buoyant density of the liposomes^54^. First, 30 μL of liposomes that incorporate the stabilized Q-SNARE complex were mixed with Nycodenz (Axis Shield, 80 %, 30 μL) and a second Nycodenz layer (30 %, 40 μL) was gently applied followed by another layer of buffer (40 μL). The density gradient was centrifuged using a Beckman TL-100 ultracentrifuge (TLS55 rotor, 100,000g, 4°C, 1 h). The 20-μL aliquots were carefully taken from the top of the gradient and analysed by coomassie blue staining.

## Statistical analysis

The statistical difference between two groups was evaluated by Student’s t-test using GraphPad Prism. Probabilities of *p* < 0.05 were considered significant.

